# Postinhibitory excitation in motoneurons can be facilitated by hyperpolarization-activated inward currents: a simulation study

**DOI:** 10.1101/2023.09.06.556472

**Authors:** Laura Schmid, Thomas Klotz, Oliver Röhrle, Randall K. Powers, Francesco Negro, Utku Ş. Yavuz

## Abstract

Postinhibitory excitation is a transient overshoot of a neuron’s baseline firing rate following an inhibitory stimulus and can be observed *in vivo* in human motoneurons. However, the biophysical origin of this phenomenon is still unknown and both reflex pathways and intrinsic motoneuron properties have been proposed. We hypothesized that postinhibitory excitation in motoneurons can be facilitated by hyperpolarization-activated inward currents (h-currents). Using an electrical circuit model we investigated how h-currents can modulate the postinhibitory response of motoneurons. Further, we analyzed the spike trains of human motor units from the tibialis anterior muscle during reciprocal inhibition. The simulations revealed that the activation of h-currents by an inhibitory postsynaptic potential can cause a short-term increase in a motoneuron’s firing probability. This result suggests that the neuron can be excited by an inhibitory stimulus. In detail, the modulation of the firing probability depends on the time delay between the inhibitory stimulus and the previous action potential. Further, the strength of the postinhibitory excitation correlates with the amplitude of the inhibitory stimulus and is negatively correlated with the baseline firing rate as well as the level of input noise. Hallmarks of h-current activity, as identified from the modeling study, were found in 50 % of the human motor units that showed postinhibitory excitation. This study suggests that h-currents can facilitate postinhibitory excitation and act as a modulatory system to increase motoneuron excitability after a strong inhibition.

**Author Summary:** Human movement is determined by the activity of specialized nerve cells, the motoneurons. Each motoneuron activates a specific set of muscle fibers. The functional unit consisting of a neuron and muscle fibers is called a motor unit. The activity of motoneurons can be observed noninvasively in living humans by recording the electrical activity of the motor units using the electromyogram. We studied the behavior of human motor units in an inhibitory reflex pathway and found an unexpected response pattern: a rebound-like excitation following the inhibition. This has occasionally been reported for human motor units, but its origin has never been systematically studied. In non-human cells of the neural system, earlier studies reported that a specific membrane protein, the so-called h-channel, can cause postinhibitory excitation. In our study, we use a computational motoneuron model to investigate whether h-channels can cause postinhibitory excitation as observed in the experimental recordings. Using the model, we developed a method to detect features of h-channel activity in human recordings. Because we found these features in a half of the recorded motor units, we conclude that h-channels can facilitate postinhibitory excitation in human motoneurons.

## 1 Introduction

Sherrington noted in his 1909 studies that “reflex inhibition of the vastocrureus and other extensor muscles in decerebrate rigidity is followed regularly, under certain circumstances, on withdrawal of the inhibitory stimulus, by a rebound contraction” [1]. Interestingly, the “rebound contraction” persisted after de-afferentation [1]. Later, the same phenomenon of motor unit postinhibitory excitation was reported in several studies [e.g., 2, 3, 4, 5, 6]. The proposed mechanisms include reflex pathways activated by muscle spindles as well as intrinsic neuron characteristics originating from the behavior of specific ion channels, but none of these studies has systematically investigated the origin of the phenomenon.

Similar to Sherrington’s findings, we observed a consistent rebound-like excitation following electrically evoked reciprocal inhibition in human tibialis anterior motor units [7] (Fig 1c). Reciprocal inhibition is a component of the stretch reflex pathway, which is disynaptically elicited by muscle spindles of an antagonistic muscle [8, 9] (Fig 1a).

**Fig 1.**
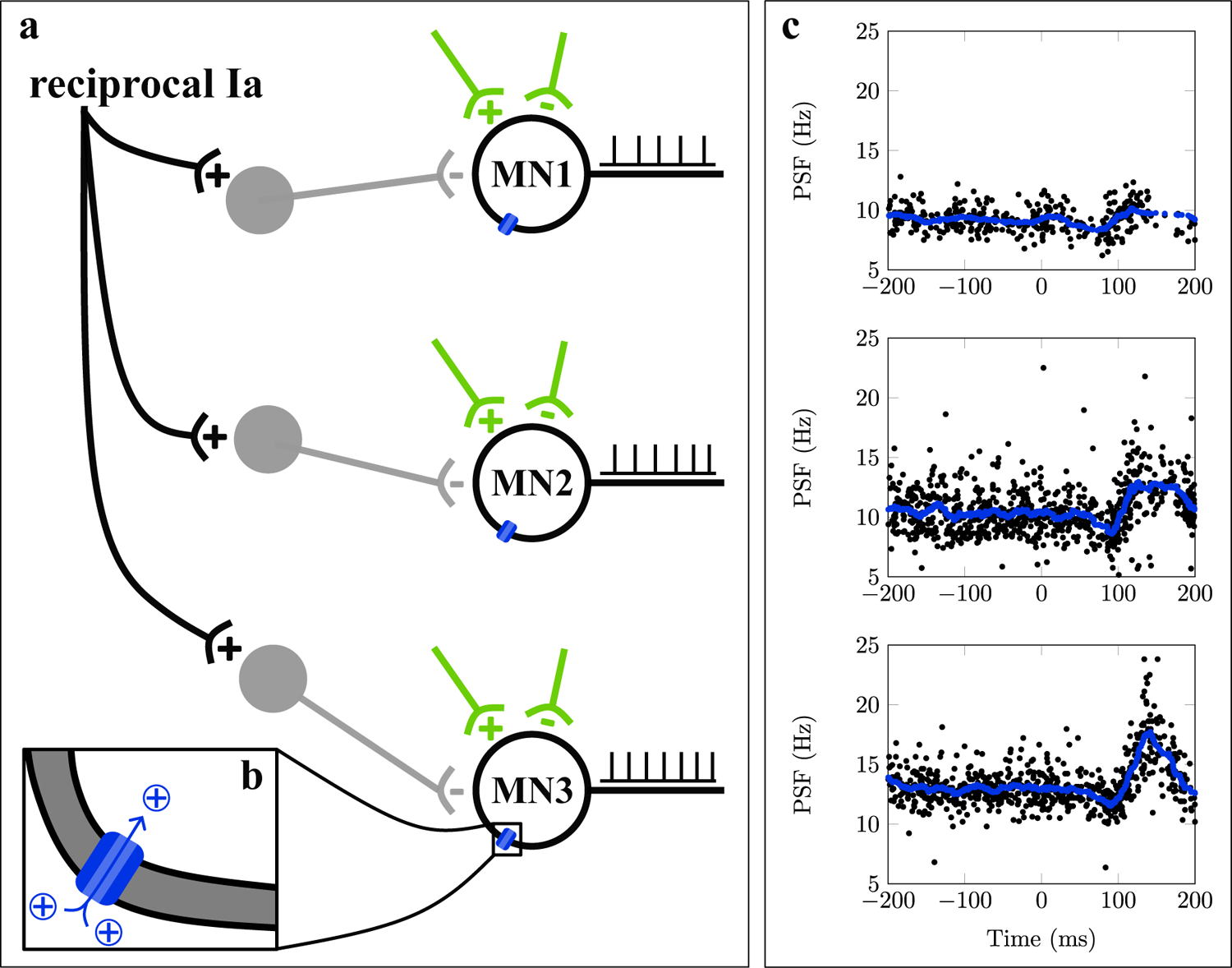
Postinhibitory excitation in human motor units. (a): Motoneurons (MNs) receive multiple, typically unknown synaptic inputs (green). Particularly, reciprocal inhibition is mediated by interneurons (gray). In living humans, the recording of motoneuron spike trains (black) allows for studying the function of the neuromuscular system. (b): The integration of synaptic inputs in motoneurons is determined by ion channels (blue). (c): Peristimulus frequencygram (PSF) of three exemplary motor units in response to reciprocal inhibition. Moving average in blue, data from Yavuz et al. [7]. It is unclear if the observed posinhibitory excitation is caused by neural pathways or intrinsic motoneuron properties.

Increased excitability elicited by a purely inhibitory stimulus is a well-known phenomenon in different cells of the mammalian neural system and a number of different ion channels was found to cause or facilitate this behavior *in vitro* [e.g., 10, 11, 12, 13, 14]. Therefore, it is surprising that intrinsic motoneuron mechanisms are rarely considered to explain the postinhibitory increase in activity of motor units *in vivo*. Instead, excitatory synaptic inputs based on reflex pathways were lately considered [e.g., 2, 3, 5, 6].

Ito and Oshima [11] first described membrane potential overshoots caused by hyperpolarization-activated inward currents in cat spinal motoneurons. Takahashi [15] found that this current, which is commonly named h-current or I_h_, is mediated by sodium and potassium ions. The corresponding channel family was identified as hyperpolarization-activated cyclic nucleotide-gated non-selective cation channels (HCN channels), which are expressed throughout the soma and dendrites of spinal motoneurons [16]. Here, we refer to this group of channels as h-channels and the corresponding currents as h-currents.

Based on the large body of evidence for intrinsic postinhibitory excitation mechanisms from *in vitro* studies, we hypothesized that intrinsic motoneuron mechanisms, specifically h-currents, also contribute to the postinhibitory excitation in motor units observed *in vivo* (Fig 1b). In this study, we analyzed for the first time the incidence and amplitude of postinhibitory excitation in human motor units following electrically elicited reciprocal inhibition of the tibialis anterior muscle. Then we used a computational motoneuron model to reproduce the experimental results and understand the role of h-currents in postinhibitory excitation. To this end, we employed a compartmental electric circuit model, which is based on previous works by Cisi and Kohn [17], Negro and Farina [18] and Powers et al. [19]. We analyzed and compared the discharge behavior of both *in silico* and *in vivo* motoneurons using the peristimulus frequencygram (PSF) [20].

## 2 Results

First, we will describe the postinhibitory excitation pattern observed in human motor units. Then, we will analyze the conditions under which we observed postinhibitory excitation in the simulated motoneuron. In the simulated motoneuron, we will determine the hallmarks of h-current mediated postinhibitory excitation. The insights gained will be used to analyze the experimental data for possible evidence of hyperpolarization-activated inward current activity.

### 2.1 Postinhibitory excitation in human motor units

We examined the incidence and amplitude of postinhibitory excitation in 159 motor units identified during the reciprocal inhibition experiment. Therefor, we analyzed the peristimulus frequencygram (PSF), which shows the instantaneous discharge rates of motor units relative to the stimulus time [20]. The amplitude of inhibition and excitation responses was determined from the cumulative summation of PSF (PSF-cusum) as described in Section 4.5. PSF and PSF-cusum of an exemplary motor unit are shown in Fig 2. In the prestimulus period, the PSF-cusum is characterized by oscillations around zero (Fig 2b). The inhibitory response is indicated by a persistent drop in PSF-cusum following the stimulus, which was applied at time zero. Notably, the time delay between the electric stimulus and the onset of the inhibition is caused by the axonal action potential conduction as well the intrinsic motoneuron latency. Postinhibitory excitation can be recognized by a subsequent sustained increase in PSF-cusum.

**Fig 2.**
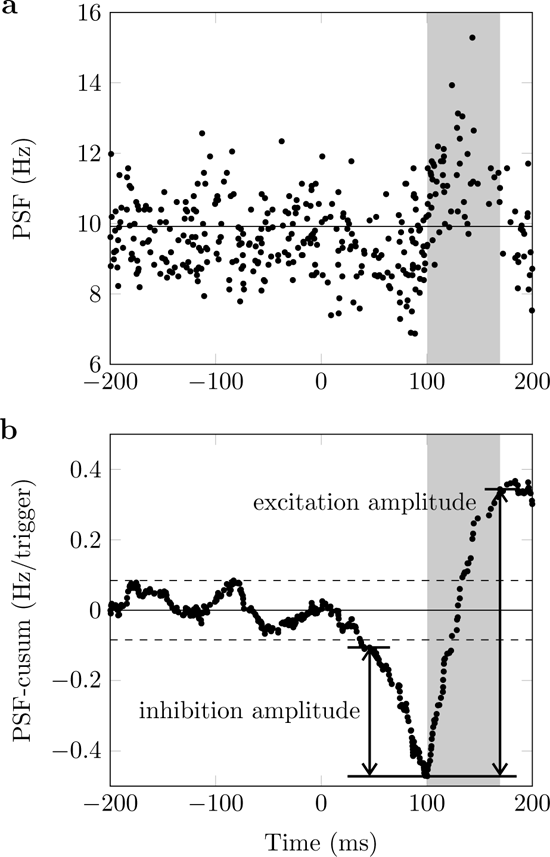
Peristimulus analysis for an exemplary selected motor unit. (a): Peristimulus frequencygram (PSF). (b): Cumulative summation of PSF (PSF-cusum). The electrical stimulus was applied at time zero. Solid horizontal lines show prestimulus mean values and dashed lines mark the significance threshold for reflex responses. Arrows show the distance between two turning points in PSF-cusum, i.e., inhibition and excitation amplitude. The period of postinhibitory excitation is highlighted with gray color. Data from Yavuz et al. [7].

All examined motor units had a significant inhibition response and the mean inhibition amplitude was 0.67 *±* 0.47 Hz*/*trigger. In total, 89 of 159 motor units showed significant postinhibitory excitation. The mean amplitude of the postinhibitory excitation was 1.80 *±* 2.08 Hz*/*trigger. The mean baseline discharge rate of the analyzed motor units was 10.12 *±* 1.66 Hz.

### 2.2 Postinhibitory excitation in simulated neurons

The results for one exemplary reciprocal inhibition simulation are summarized in Fig 3. Thereby, we directly compared the behavior of the motoneuron model with h-current (Fig 3a) and the h-current knock-out model (Fig 3b). Before the stimulus, both simulated neurons show a stationary baseline activity, i.e., stable mean discharge rate and interspike interval variability. After a silent period both motoneurons show an inhibitory response, which is apparent by a decrease in PSF-cusum (Fig 3c, d). Although the same inhibitory stimulus was applied to both computational neurons, the inhibition amplitude is larger in the neuron without h-current (1.11 Hz*/*trigger vs. 0.34 Hz*/*trigger). After the inhibition only the neuron with h-current shows a significant excitation response (amplitude 0.33 Hz*/*trigger).

**Fig 3.**
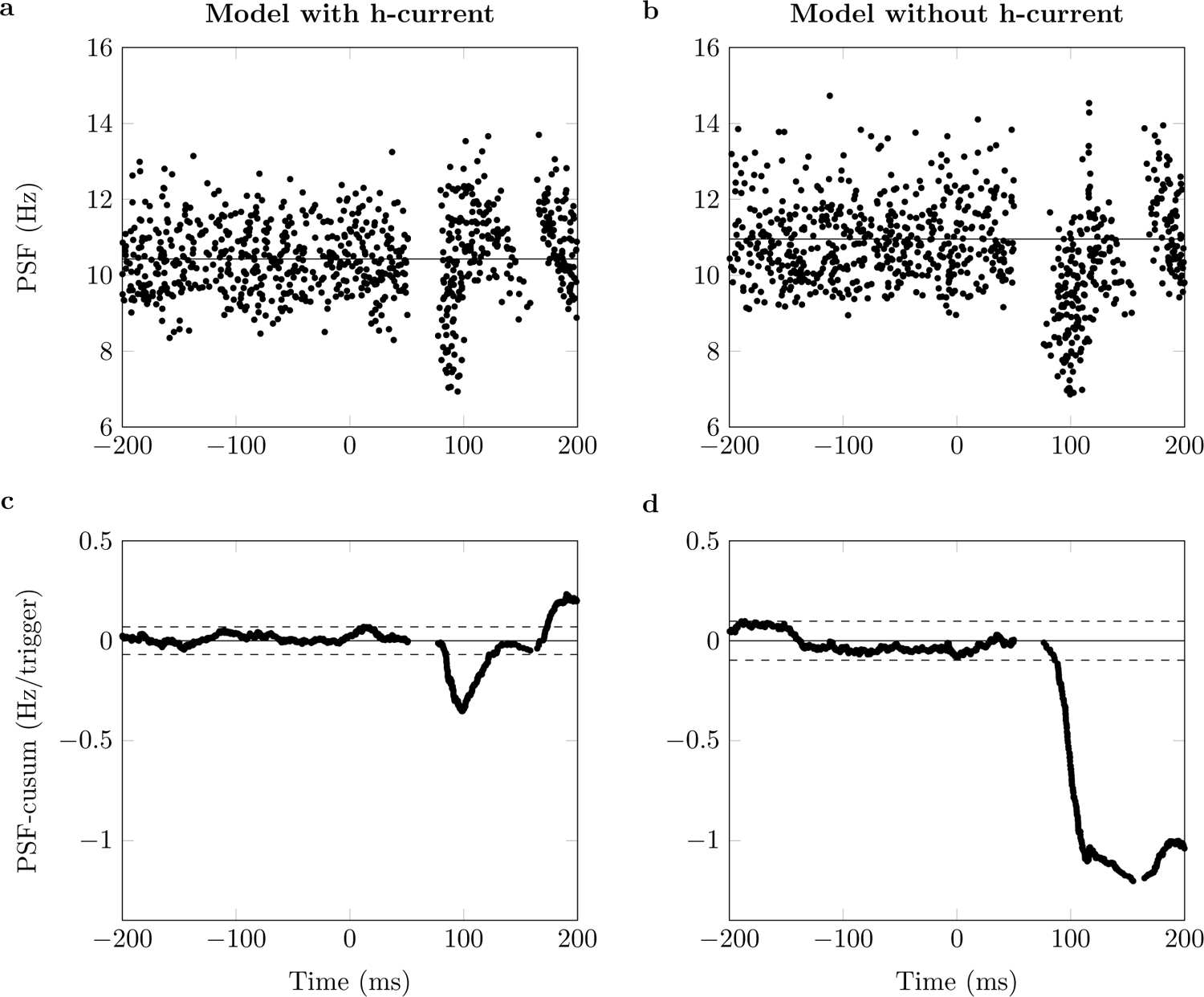
Peristimulus analysis for simulated motoneurons. (a): Peristimulus frequencygram (PSF) for a simulated neuron with h-current. (b): PSF cumulative summation (PSF-cusum) for a simulated neuron with h-current. (c): PSF for a simulated neuron without h-current. (d): PSF-cusum for a simulated neuron without h-current. The stimulus was applied at time zero. Solid horizontal lines show prestimulus mean values and dashed lines mark the significance threshold for reflex responses. Inhibitory stimulus amplitude *−*10 nA.

To investigate the role of different factors in postinhibitory excitation in more detail, we simulated reciprocal inhibition of the motoneuron with h-current under various conditions, i.e., varying the inhibitory stimulus amplitude, the mean drive and the input noise level. This yielded baseline frequencies between 9.99 Hz and 18.52 Hz and a coefficient of variation of the interspike interval ranging from 0% to 12 % (Table 1). Notably, it is observed that increasing the noise also yields a higher baseline discharge rate. This can be explained by the fact that at high noise random oscillations of the membrane potential are more likely to hit the depolarization threshold.

**Table 1.** Results of the linear regression analysis. Mean frequency and coefficient of variation of the interspike interval (CoV ISI) of the baseline activity as well as slope and coefficient of determination (R^2^) values of the linear regression for nine different combinations of mean drive and noise input are given. Linear regression was performed using the least-squares method, provided that more than one data point was available (otherwise marked with n.a.)

| Noise | Drive | Baseline frequency (Hz) | CoV ISI (%) | Slope (Hz/trigger/nA) | $R^2$ |
| --- | --- | --- | --- | --- | --- |
| No | Low | 9.994 | 0.048 | 0.042 | 0.998 |
|  | Medium | 13.995 | 0.069 | 0.023 | 0.997 |
|  | High | 17.994 | 0.079 | 0.013 | 0.993 |
| Low | Low | 10.431 | 8.718 | 0.03 | 0.981 |
|  | Medium | 14.153 | 6.395 | 0.02 | 0.999 |
|  | High | 18.026 | 5.98 | n.a. | n.a. |
| High | Low | 11.255 | 12.123 | 0.011 | 0.983 |
|  | Medium | 14.602 | 10.166 | n.a. | n.a. |
|  | High | 18.53 | 10.108 | n.a. | n.a. |

For all conditions, the simulated motorneuron showed an inhibition response with inhibition amplitudes ranging from 0.26 Hz*/*trigger to 1.67 Hz*/*trigger. Yet, an excitation response was not always observable. The simulated motoneuron showed postinhibitory excitation amplitudes between 0 Hz*/*trigger, i.e., no significant excitation response, and 0.64 Hz*/*trigger. Fig 4 summarizes the results from all simulations. It can be observed that the excitation amplitude increases with the size of the inhibitory stimulus. When fixing both the mean drive and the noise level, the relationship between the inhibitory stimulus size and the excitation amplitude can be approximated with a linear regression model (98% *<* R^2^ *≤* 99.9%, Table 1). For a fixed stimulus size, the excitation amplitude, as well as the slope of the linear regression, are higher for a lower drive, i.e., smaller baseline firing rates (Table 1). Moreover, the amplitude of the postinhibitory excitation is negatively correlated with the amount of noise. At high noise and high drive, no excitation response can be observed at all.

**Fig 4.**
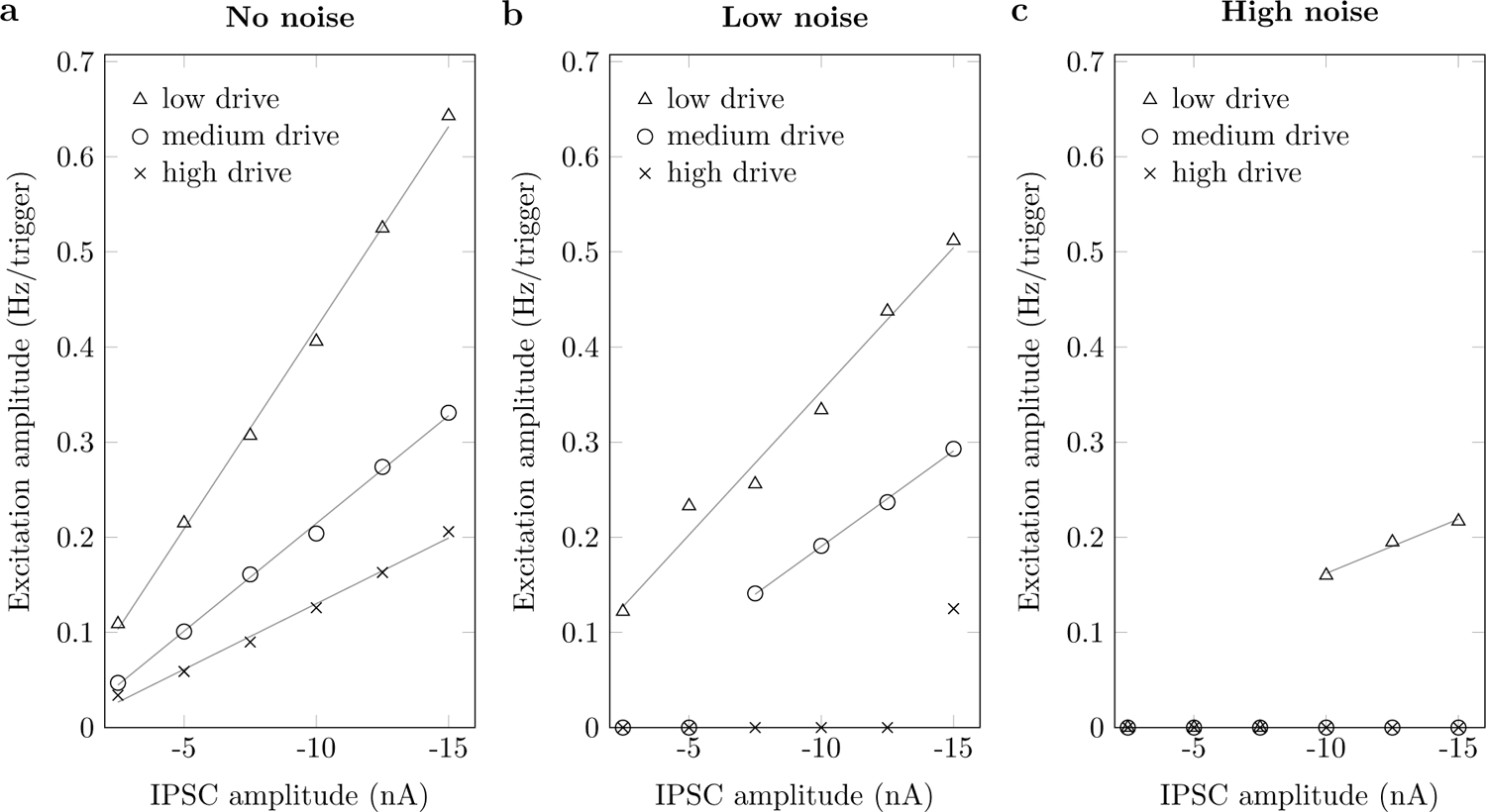
Postinhibitory excitation in different simulation settings. Excitation amplitudes in relation to inhibitory postsynaptic current (IPSC) amplitude and for three different baseline discharge rates (low (*△*), medium (o) and high (x) drive) and with three different amounts of noise (standard deviation 0 % (a), 12.5 % (b) and 25 % (c) of mean drive). Gray lines show linear regressions.

In summary, postinhibitory excitation could only be observed in simulated neurons with hyperpolarization-activated inward currents. Systematically varying the model parameters revealed that the postinhibitory excitation amplitude is correlated with the strength of the inhibitory stimulus. Yet, the excitation amplitude is modulated by the mean discharge rate and the level of noise. That is, increasing the baseline activity or noise decreases the excitation amplitude, even to a degree where excitation is no longer observed.

### 2.3 Post-inhibitory excitation is stimulus-time dependent

We made use of the *in silico model* to mechanistically link the observed postinhibitory excitation responses and the biophysical behavior of motorneurons. This was possible because the simulation allows to observe all system parameters and relate them to each other. The effect of hyperpolarization-activated inward currents was isolated by comparing the trajectories of the membrane potential of computational motoneurons with and without h-channel. Here, we considered an exemplary simulation with low drive and an inhibitory stimulus amplitude of *−*10 nA. To focus on the effect of the h-current, we also omitted noise in these simulations.

The simulations revealed that when applying an inhibitory current pulse the instantaneous discharge rate of the simulated motoneurons always depends on the time delay between the stimulus and the previous motoneuron discharge t_stim_ (Fig 5). Particularly, the model with h-current showed two opposing responses (Fig 5a, b). For large t_stim_ values, the next action potential was delayed compared to an unperturbed reference simulation, i.e., the neuron was inhibited. In contrast, for a shorter t_stim_, the interspike interval was shortened compared to the undisturbed reference simulation, i.e., the neuron was excited. Thereby, the amount of excitation depends on the exact timing of the stimulus. The insert in Fig 5a shows that the inhibitory stimulus induced a positive inward ionic current flux that continued even after the end of the stimulus. Thus, the additional depolarization (compared to the unperturbed reference simulation shown in black color) is caused by the h-current. Excitation in response to inhibitory stimuli was observed only in simulations with h-current. In the model without h-current, the inhibitory stimulus always inhibited the neuron. Thereby, later stimuli (large t_stim_) caused a stronger inhibition than earlier stimuli (Fig 5c, d).

**Fig 5.**
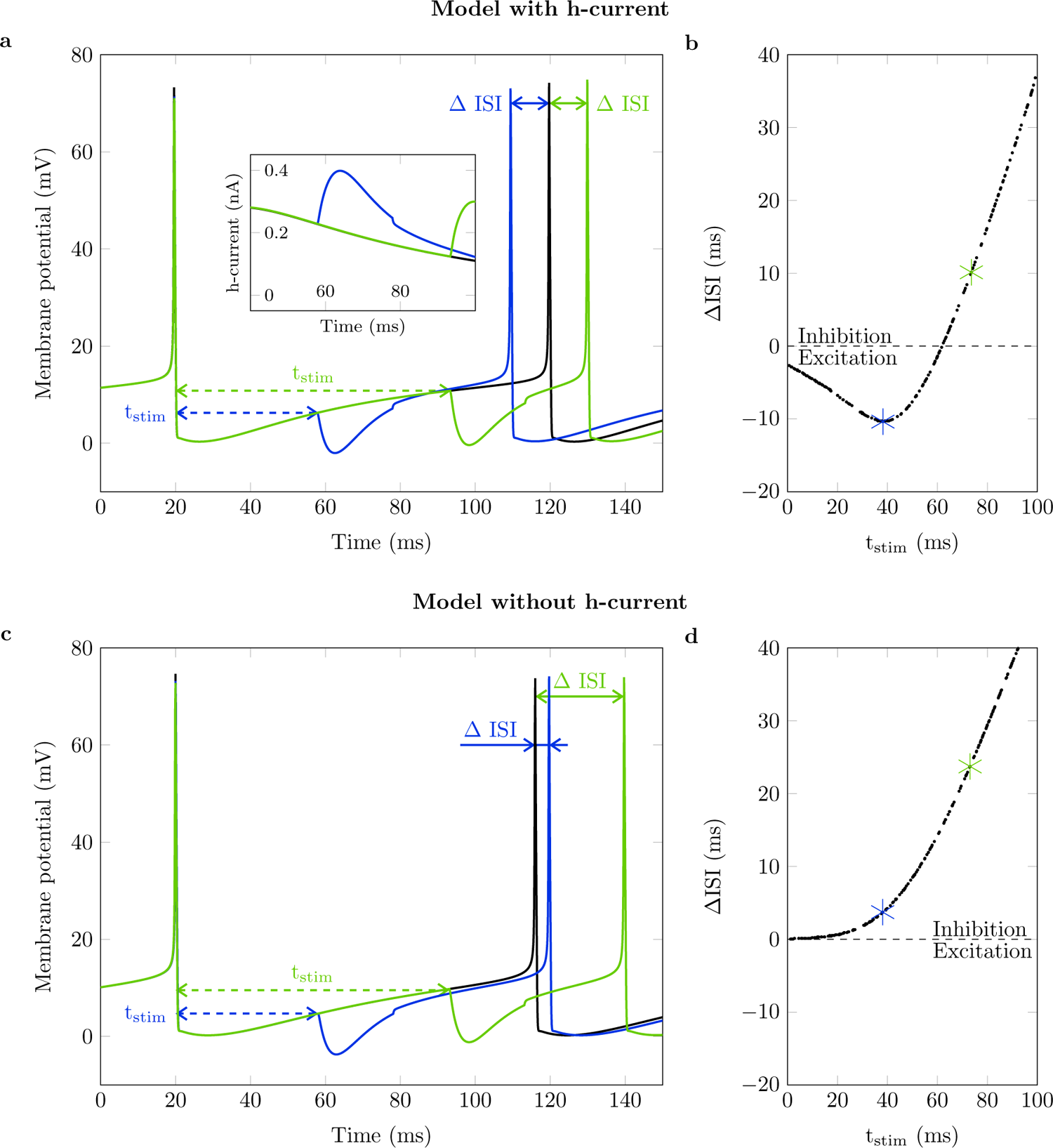
Analysis of stimulus time-dependent interspike interval duration in simulated motoneurons. Top row: model with h-currents, bottom row: model without h-currents. Baseline frequency 10 Hz, no noise, inhibitory stimulus amplitude *−*10 nA. (a, c): Membrane potential trajectory without stimulus (black, undisturbed interval) and with inhibitory stimulus applied at two exemplary time points (blue, green). Continued on next page. Dashed arrows mark inhibitory stimulus time with respect to last discharge (t_stim_) and solid arrows mark change of interspike interval with respect to the undisturbed interval (Δ ISI). Insert in (a) shows h-current for the shown interspike intervals between 40 ms and 100 ms. Here, positive sign indicates current flux into the cell. (b, d): Change of interspike interval duration (Δ ISI) over time of stimulus application with respect to the last discharge (t_stim_). Intervals shown in (a) and (c), respectively, are marked with asterisks. Dashed lines separate prolonged interspike intervals (inhibition) from shortened interspike intervals (excitation).

This characteristic opposing behavior of the computational motoneurons with hcurrent can be also visualized in the previously shown PSF and PSF-cusum plots. Therefore, the spike trains shown in the PSF plots were clustered into two groups. In detail, we separated spike trains where the instantaneous discharge frequency of the first poststimulus spike was significantly (by more than one standard deviation) higher or lower than the baseline discharge rate. Fig 6 shows the same data as previously reported in Fig 2 and Fig 3. However, spike trains where the first postinhibitory spike shows significant excitation are visualized in blue color (excitation cluster) and spike trains where the first postinhibitory spike shows significant inhibition are shown in green color (inhibition cluster). The results of an exemplary simulated motoneuron with h-current is shown in Fig 6b, e and clearly shows distinct excitation and inhibition clusters. This is also evident from the PSF-cusums of the clustered spike trains (Fig 6g). The initial decrease in overall PSF-cusum (black) is caused by the inhibition cluster (green). As expected, the short latency after the stimulus as well as the timing of the prestimulus spikes indicate that the stimulus was delivered late in the motoneuron’s integration phase (large t_stim_, cf. Fig 5). The initial inhibition is followed by the excitation response (blue) that ultimately predominates the overall PSF-cusum. The longer latency of the spikes associated with the excitation cluster as well as the pattern of the prestimulus spikes are in agreement with our finding that excitation is observed when the stimulus is delivered early in the integration phase of the motoneuron (small t_stim_). In conclusion, it is possible to find evidence for hyperpolarization-activated inward current activity by means of PSF analysis.

**Fig 6.**
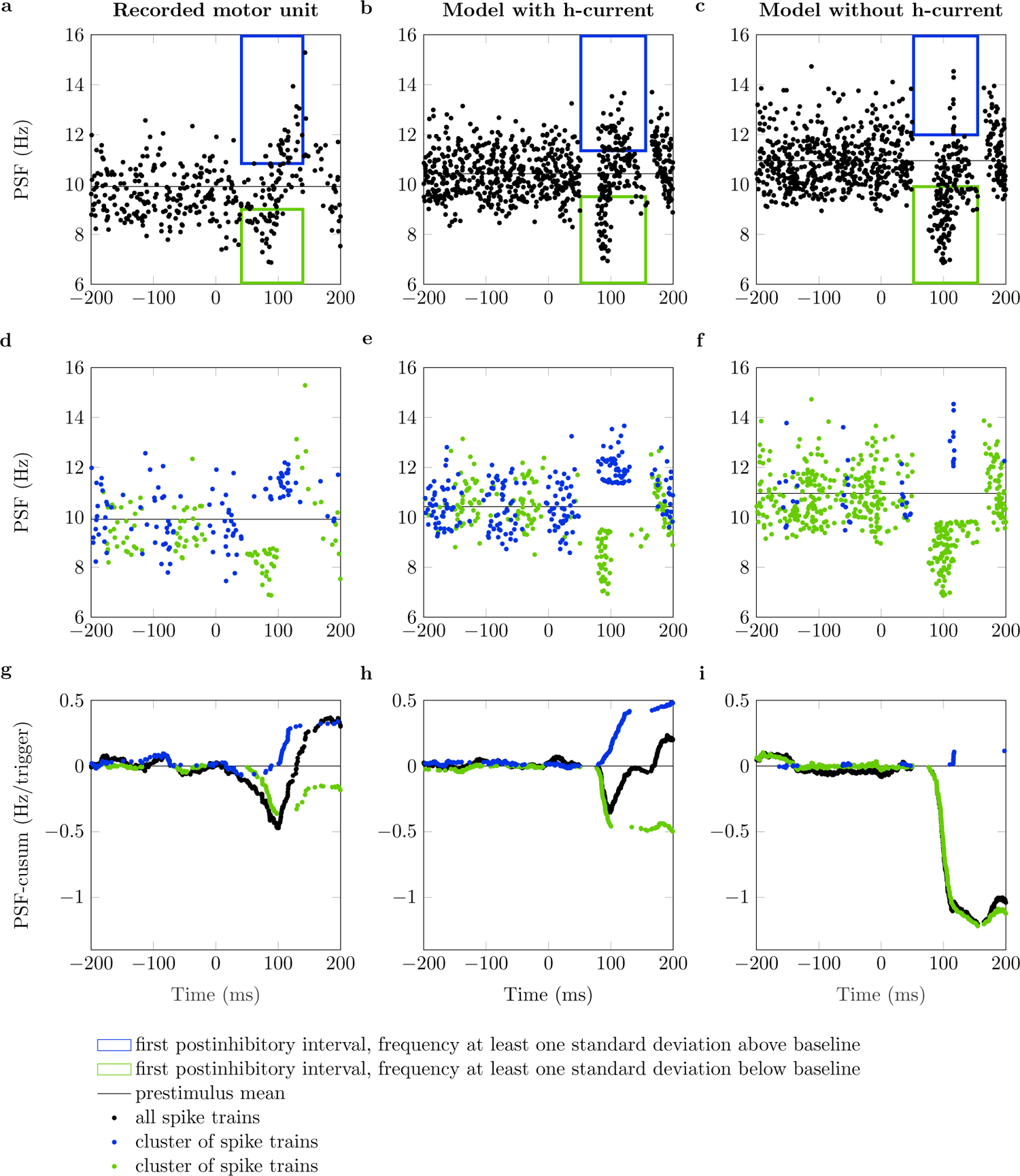
Cluster-based analysis of peristimulus frequencygram. PSF and PSF cumulative sum (PSF-cusum) for one experimentally recorded motor unit and a simulated neuron with and without h-current (data from Fig 2 and Fig 3). The blue boxes in panels (a, b, c) cluster the first postinhibitory spikes that fire at least one standard deviation above the mean baseline frequency (black line). Accordingly, the green boxes cluster all first postinhibitory spikes that fire with at least one standard deviation below the mean baseline frequency. (d, e, f): Clusters of spike trains from which the first poststimulus spikes appear in the blue or green box, respectively. (g, h, i): PSF-cusum of all spike trains (black) and clusters of spike trains (blue, green).

Hence, we investigated experimentally recorded motor units that showed significant postinhibitory excitation with the described cluster-based PSF analysis. In total, 45 of 89 motor units with significant postinhibitory excitation showed frequencies significantly higher than the baseline frequency in the first poststimulus interval (to at least 10 % of delivered stimuli). The results for an exemplary chosen experimentally recorded motor unit are shown in Fig 6a, d, h. Indeed, one can observe the same characteristic behavior as for the simulated motoneuron with h-current, i.e., distinct inhibition and excitation clusters, whereby the inhibition precedes the excitation.

For completeness, the analysis was also performed for the simulated neuron without h-current. As expected, for the neuron without h-current the number of samples in the excitation cluster is very low and does not cause a relevant excitation response (Fig 6c, f, i).

From the analysis of the membrane potential and PSF of the simulated motoneurons, it can be concluded that h-currents cause the motoneuron to be either inhibited or excited following an inhibitory stimulus, depending on the timing of the stimulus with respect to the last discharge. This characteristic response pattern was also found in the first poststimulus interval of half of the recorded motor units.

## 3 Discussion

We identified an excitatory response that frequently follows a strong reciprocal inhibition in motor units of the human tibialis anterior muscle. For the first time in motor units recorded *in vivo*, we investigated if an intrinsic motoneuron property can cause this phenomenon. Particularly, we hypothesized that a hyperpolarization-activated inward current (h-current) may be one of the factors that contribute to this excitatory response pattern. Using a computational motoneuron model, we showed that the h-current could lead to an excitation response after an inhibitory stimulus when; 1) the inhibitory stimulus is applied at an appropriate time window in the neuron’s integration phase, 2) the inhibitory stimulus has a sufficiently large amplitude and 3) other inputs to the neuron are small. Further, it is was shown that evidence for hyperpolarization-activated inward current mediated postinhibitory excitation can be tested *in vivo* by means of cluster-based PSF analysis. The integrated evaluation of both *in vivo* and *in silico* data presented within this study shows that h-currents could be a mechanism that facilitates postinhibitory excitation in motoneurons. This challenges an established paradigm that postinhibitory excitation in motoneurons is dominantly caused by reflex pathways and underlines that intrinsic motoneuron properties have to be considered when interpreting *in vivo* reflex experiments.

### 3.1 Insights on mechanisms of postinhibitory excitation

A detailed analysis of the *in silico* experiments showed that h-currents cause a non-linear, history-dependent input-output relation for short inhibitory stimuli. That is, excitation or inhibition depending on the delay between the inhibitory stimulus and the previous motorneuron discharge. H-currents are activated at hyperpolarized membrane potentials [e.g., 21]. Counterintuitively, an inhibitory stimulus can therefore lead to an increased net current influx into the cell. If the inhibitory stimulus is applied when the motorneuron is close to its depolarization threshold, the instantaneous effect of the inhibitory stimulus is dominant. Hence, one will observe inhibition. However, if the motorneuron is inhibited earlier in the integration phase, the stimulus-induced additional h-current influx ultimately dominates the inhibitory stimulus and causes excitation.

The opposing motorneuron responses can be visualized for both *in silico* and *in vivo* data through a cluster-based PSF analysis. Thereby, spike trains are grouped according to the relative instantaneous discharge rate of the first poststimulus discharge. It was shown that a considerable fraction of the experimentally recorded motor units showed characteristic clusters of both excitation and inhibition. The same patterns were observed for the simulated motorneurons with h-currents. Hence, these finding provide evidence for h-current activity in human motor units.

The model predicted that the excitation response is negatively correlated with a motorneuron’s firing rate and can even become undetectable at high noise levels. In this work, the ‘high noise’ condition was characterized by a coefficient of variation of the interspike interval ranging from 10 % to 12 %. This is at the lower end of what was reported for humans [e.g., 22, 23]. This potentially explains our finding that the excitation response was not observable in all investigated motor units. Long inhibitory stimuli in combination with high discharge rates might have also prevented Türker and Powers [4] from observing excitation in the first poststimulus interval when they used a comparable cluster-based PSF analysis to investigate the effect of large inhibitory postsynaptic potentials in rat hypoglossal motoneurons.

Notably, also other factors may modulate h-current mediated postinhibitory excitation. For example, persistent inward currents (PICs) can modulate h-current activity [24]. However, PICs were shown to play a minor role during Ia reciprocal inhibition [25]. Further, due to their slow time dynamics, we expect no major changes in PIC activity during short inhibitory stimuli [26]. Importantly, modulations in h-current conductance have no influence on the characteristic opposing motorneuron response (Fig S 2b).

Postinhibitory excitation of motorneurons is a well-known phenomenon that was repeatedly observed both in living humans and for *in vitro* preparations [e.g., 4, 5, 6]. Yet, no consent regarding the biophysical origin of this phenomenon has been firmly established. Previously proposed mechanisms include intrinsic motorneuron properties like summation effects of the ionic channel conductances that are active during the afterhyperpolarization [4] or neuronal pathways, e.g., Ia and II stretch reflexes [5, 6]. In the presented simulations, we did not consider other neuronal pathways that potentially contribute to the postinhibitory excitation. Importantly, the results shown in this work only consider the first poststimulus discharge. On this time scale, the involvement of a neural pathway is unlikely. Regarding other internal mechanisms, in the simulations we could observe a mild effect of afterhyperpolarization summation that leads to increased discharge rates of the second poststimulus spike. Further, additional simulations with a longer time constant for the h-channel showed that, under certain circumstances, the h-current can last long enough to slightly increase the frequency of the second poststimulus discharge of the neuron.

We conclude that h-currents can facilitate postinhibitory excitation observed at the first poststimulus discharge [e.g., 5, 7]. However, to fully uncover the complex behavior of motoneurons in response to an inhibitory stimulus other factors need to be considered. Particularly, postinhibitory excitation at the second poststimulus discharge [e.g., 4, 6] cannot be fully explained by h-currents.

### 3.2 Limitations

In this study, we used a computer model to assist the interpretation of empirical observations from reflex experiments in living humans. However, it is currently not feasible to measure all model parameters that would be needed to directly replicate the corresponding *in vivo* experiment. Thus, the presented simulations are a simplification of the underlying physiology.

The utilized model reduces the structure of the dendritic tree into a single model compartment and does not explicitly describe all channels that can be found in human motoneurons. Yet, previous studies showed that a two-compartment model with a limited number of conductances can replicate realistic motoneuron firing patterns [17, 18]. Adding h-channels yields the simplest model that can sufficiently address the posed research question. The h-channel modulates the rheobase of the simulated motorneuron. To compare the model with h-currents and the knock-out model the input current was adjusted, i.e., both models operate at comparable points of their current-frequency relation. This guarantees that differences are exclusively attributed to h-currents (see Fig S 1 for the gating variables). The parameterization of the h-channel can reproduce the magnitude of membrane potential overshoots in response to hyperpolarizing currents steps as observed *in vitro* [12, 27]. To compensate for the temperature difference between the *in vitro* experiment and *in vivo* conditions we followed a time-dynamics correction proposed by Powers et al. [19]. Although it is impossible to directly validate the utilized implementation, additional simulations showed that varying the h-channel parameters does not affect the general behavior of the model (Fig S 2).

Reciprocal inhibition indirectly stimulates the motorneurons through the afferent nerve. The magnitude of the inhibitory stimulus entering the motoneuron through interneurons is unknown. To compare the simulations and experimental data, the size of the inhibitory stimulus was chosen such that the inhibition amplitudes determined from PSF-cusum are in the same range as for the experimental study. Nevertheless, a quantitative comparison of the postinhibitory excitation amplitudes of *in silico* and *in vivo* motoneurons is beyond the scope of this study.

Further, instead of explicitly modeling synapses the input signals represent effective synaptic currents, i.e., currents that eventually reach the soma. This is a reasonable simplification since only these currents affect the generation of action potentials [28]. Still, injecting the inhibitory stimuli into the soma bypasses h-channels located on the dendrite to a certain extent and hence, underestimates the effect of h-channels. Synapses for reciprocal inhibition are located close to the soma [29, 30, 31], and thus, the thereby introduced error is assumed to be small.

Postinhibitory excitation is a phenomenon that was repeatedly shown *in vitro* in different species and cell types and attributed to different mechanisms, e.g., h-channels, (calcium-activated) potassium conductances, low threshold sodium conductances, delayed recovery of the sodium inactivation gate (anode break excitation), T-type calcium channels, or NMDA receptors [32, 12, 13, 27, 33, 14, 34]. One particular strength of computer models is the possibility to study the influence of an isolated phenomenon. The family of h-channels is a likely candidate as its gating variable shows steep slopes for hyperpolarized membrane potentials and its time constant allows it to react at the timescale of one interspike interval [21, 35, 19]. Note that anode break excitation can be excluced in our model since the sodium inactivation gate is almost fully open at resting potential (Fig S 1). We cannot rule out the possibility that one or several of these channel mechanisms may also contribute to the postinhibitory excitation in motor units. Nevertheless, it highlights that internal motorneuron properties need to be considered for the investigation of postinhibitory excitation in human motor units.

### 3.3 Functional significance and future directions

We showed that h-current mediated postinhibitory excitation is most pronounced when a motorneuron operates close to its recruitment threshold and for strong inhibitory stimuli. Thus, h-currents temporally increase the excitability of a motoneuron and potentially protect a motor unit from derecruitment. In previous studies, h-currents were shown to increase the excitability of human motor axons after hyper-polarization and to play a role in certain diseases associated with hyperexcitability, e.g., neuropathic pain and restless legs syndrome [36, 37, 38]. Our results indicate that h-currents can act as an ultra-fast and short-term adaptation mechanism in the motor control system that fine-tunes spinal excitability.

Coupling the motoneuron model with a skeletal muscle model could provide further insights into the functional role of postinhibitory excitation and its influence on, e.g., force steadiness and force variability [e.g., 39]. Future modeling studies should further address the different possible causes of postinhibitory excitation each in isolation but in comparison to the other. Nevertheless, the most promising way to quantify the contribution of h-currents to postinhibitory excitation is *in vitro* studies, also due to the uncertainty of the channel parameters in the model. This study pointed out that it is crucial to choose evaluation and analysis methods, which can take activation history into account.

### 3.4 Summary and Conclusion

We investigated how hyperpolarization-activated inward currents (h-currents) can contribute to postinhibitory excitation in human motor units. Using a computational model we showed that h-currents can shorten interspike intervals in response to strong inhibitory stimuli and thus, facilitate postinhibitory excitation. This effect is stimulus time-dependent and most pronounced in conditions with low firing rates and low noise, i.e., few other inputs. Furthermore, this study showed that intrinsic motor neuron properties must be considered for the interpretation of reflex responses. The presented PSF cluster method reveals history-dependent effects. Excitation in the first poststimulus interval after reciprocal inhibition was found in a significant portion of the analyzed human motor units. We conclude that h-currents serve as a modulatory system that is able to increase motoneuron excitability.

## 4 Methods

This section provides an elaborate description of the utilized computational model and data analysis techniques. The experimental data analyzed in this manuscript were collected in a previous study [7] and only a brief description of the experimental protocol is provided.

### 4.1 Experimental data

Motor unit reciprocal inhibition data from a previous study [7] were used to investigate the incidence rate and strength of postinhibitory excitation activity *in vivo*. The data were acquired with the approval of the local ethical committee of the University Medical Center, Georg-August–University of Göttingen (approval date: 1/10/12). High-density surface EMG (HDsEMG) was recorded (sampling rate: 10 240 Hz) from tibialis anterior muscles during steady isometric contraction at 10 % and 20 % of the maximum voluntary contraction (MVC). Stimulating the tibial nerve through monopolar stimulation electrodes elicited reciprocal inhibition on the tibialis anterior muscle. The metal pin anode and cathode electrodes were placed on the skin of the popliteal fossa to stimulate the nerve. The activity of individual motor units was identified by decomposing HDsEMG data using a blind source separation technique [40, 41]. Further details about the experimental protocol and analysis can be found in Yavuz et al. [7].

### 4.2 Computational modeling

A single motoneuron was simulated using an electric circuit model, which is based on the motoneuron model proposed in Negro and Farina [18] and [39]. In short, the *in silico* motoneuron consists of two compartments, i.e., the soma and a lumped dendrite. The soma compartment includes voltage-gated conductances that describe sodium as well as slow and fast potassium channels. Further, both compartments include an additional leakage conductance. For the present investigation, we added a voltage-gated h-channel conductance in both the soma and the dendrite compartment (Fig 7d). Accordingly, the membrane potential in each compartment can be described by the following differential equations:

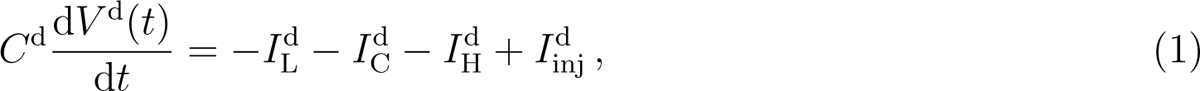

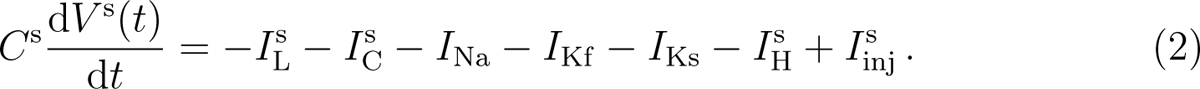

**Fig 7.**
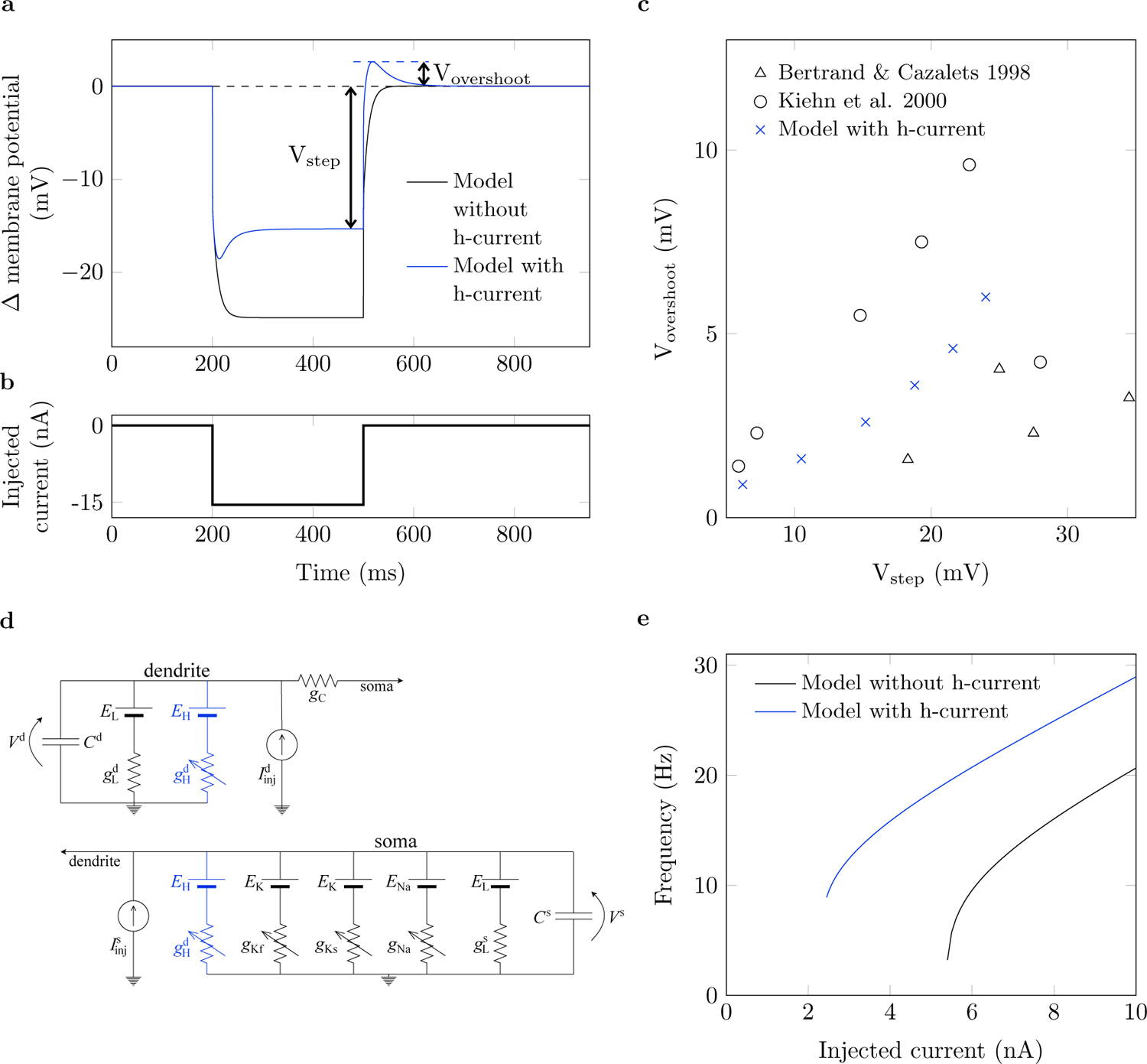
Characteristic behavior of the computational motoneuron model. (a): Membrane potential time course of the simulated motoneurons without (black) and with h-current (blue) in response to injection of the current step shown in (b). Membrane potential is given with respect to the resting potential. (c): Steady-state membrane potential (*V*_step_) vs. overshoot membrane potential (*V*_overshoot_) of the simulated neuron (x) compared to data obtained from Kiehn et al. [27] (o) and Bertrand and Cazalets [12] (*△*). (d): Equivalent electric circuit of the motoneuron model. Blue color highlights components that were added with respect to the previous version of the model [18]. (e): Current-frequency relation for the model without (black) and with (blue) h-currents [35, 21]. Note that all potentials are given with respect to the resting potential.

Therein, *V* (*t*) is the membrane potential at time *t* and *C* is the membrane capacitance. The superscripts ‘s’ and ‘d’ denote the soma and the dendrite compartment, respectively. The currents *I*_Na_*, I*_Kf_ and *I*_Ks_ describe the flux of ions through sodium and fast and slow potassium channels, respectively. Further, *I*_C_ describes the coupling current between the two compartments whereby *I*^d^ = *−I*^s^. *I*_L_ is a leakage current and *I*_inj_ denotes currents injected into the compartments, e.g., by an external electrode. Currents are modeled and parameterized as described in Negro and Farina [18] and Röhrle et al. [39]. H-channels were found to be expressed widely across motoneurons, consequently, they are placed in both soma and dendrite compartments [16]. The mathematical description of the h-current and its parameters are provided in Equations (3) to (5).

The h-current, *I*_H_, is described by Powers et al. [19]:

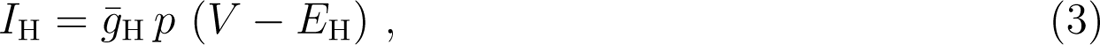

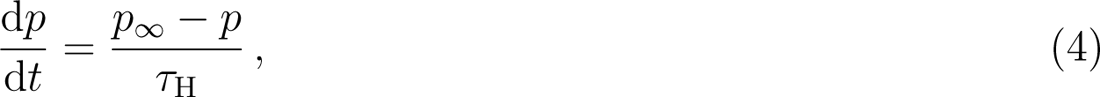

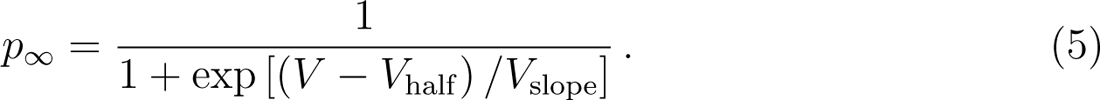

Thereby, *E*_H_ denotes the reversal potential of the h-conductance and *g*^-^_H_ the maximum conductance. The transient behavior of the gating variable *p* is determined by the voltage-independent time constant *τ*_H_, the half maximum activation potential *V*_half_ and a slope factor *V*_slope_. Note that Equations (3) to (5) refer to both compartments, i.e., *V* corresponds to *V* ^s^ and *p* to *p*^s^ for the soma compartment and to *V* ^d^ and *p*^d^ for the dendrite compartment.

### 4.3 Model parameterization

Following a study by Duchateau and Enoka [42], we assumed that for tibialis anterior muscle and contraction strengths of less than 20 % MVC, all recruited and consequently all recorded motor units are of slow type. Model parameters for different motor unit types were published by Cisi and Kohn [17] (see their Table 2). We chose the input resistance and the size parameters (radius and length) of both the soma as well as the dendrite compartment according to the mean values for slow-type motoneurons.

Since the h-channel was not considered by either Cisi and Kohn [17] or Negro and Farina [18], we used the implementation by Powers et al. [19] as a guide. Powers et al. [19] used experimental recordings by Larkman and Kelly [21] for the parameterization of the channel and we adopted the time constant as well as the slope factor, i.e., *τ*_H_ = 50 ms and *V*_slope_ = 8 mV. In contrast to Powers et al. [19], we assumed that persistent inward currents play a minor role in our study [25]. Thus, we adapted the reversal potential *E*_H_ = 20 mV and half maximum activation potential *V*_half_ = *−*20 mV with respect to the original implementation and in accordance with experimental data

The above parameters determine the temporal dynamics of the h-current. The maximum conductance *g*^-^_H_, which can also be interpreted as the density of h-channels in the membrane, determines the maximum amount of h-current that can flow across the membrane. Experimental current clamp studies from Bertrand and Cazalets [12] and Kiehn et al. [27] were used to parameterize *g*^-^_H_. Therefore, the simulated membrane potential trajectory in response to injection of hyperpolarizing current steps was compared to *in vitro* results. The simulated neuron shows the characteristic undershoot and overshoot at the beginning and end of the applied current step (Fig 7a, b), which is typically attributed to h-currents [e.g., 35, 12, 11, 21].

For a quantitative comparison, the steady-state membrane potential just before the release of the current step, *V*_step_, was compared to the maximum size of the membrane potential overshoot after the release of the current step, *V*_overshoot_ (Fig 7a, b). The current step was applied for 300 ms to reach a steady membrane potential. With a maximum conductance of *g*^-^_H_ = 2 mS cm*^−^*^2^ the simulated neuron showed overshoot potentials of 0.84 mV to 6 mV in response to current injections that hyperpolarize the membrane potential by 6 mV to 24 mV. This is well within the range of values reported in Bertrand and Cazalets [12] and Kiehn et al. [27], i.e., overshoot potentials of 0.96 mV to 9.6 mV for potential steps of 5.9 mV to 34.5 mV (Fig 7c). Note that adding the h-current does not lead to subthreshold oscillations and thus preserves the integrator nature of the motoneuron model (Fig 7a).

The above parameterization results in a total h-conductance of 25 nS (obtained from multiplying *g*^-^_H_ with the total membrane area of the simulated neuron). According to Larkman and Kelly [21] and Kjaerulff and Kiehn [43] this is within the range of experimental data (12 nS to 89 nS).

Adding the h-current increases the resting potential of the neuron due to an additional inflow of current, which was also reported by Powers et al. [19]. Consequently, the h-current model has a decreased rheobase, but the gain of the current-frequency relation is hardly affected (Fig 7e). By considering the simulated neurons at the same working point, defined by the frequency, we ensure comparability between the models.

### 4.4 Simulation of reciprocal inhibition

The applied simulation protocol aims to mimic the experimental procedure described in Section 4.1. Repeated injections of inhibitory current pulses imitate the stimulus delivered in the reflex experiment. The stimuli were applied 200 times with a random interstimulus interval of 1000*±*100 ms. All simulations were performed with MATLAB R2021a (9.10.0.2015706). The motoneuron model is represented by a system of eight ordinary differential equations, which was solved with MATLAB’s ode23 (single-step, explicit Runge-Kutta solver [44]) and an absolute and relative error tolerance of 1 *×* 10*^−^*^5^.

The activity of the motoneuron model is driven by an injected current. To replicate the experimental protocol, the input is composed of up to three components: (i) inhibitory current pulses simulating reciprocal inhibition, (ii) a constant current representing the mean cortical drive to the neuron and which determines the contraction strength, (iii) additive noise representing all afferent and efferent inputs to the motoneuron that are not explicitly modeled.

Reciprocal inhibition (i) was simulated by injection of a current kernel representing the compound inhibitory postsynaptic current (PSC). The postsynaptic current *I*_PSC_ is described by

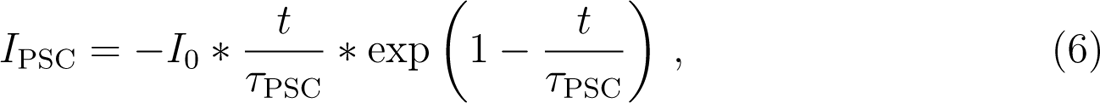

with *t* representing the time since the beginning of the current injection. The time constant *τ*_PSC_ = 4 ms was fixed [45], while the amplitude *I*_0_ ranged from 2.5 to 15 nA to cover a large range of inhibitory stimulus strengths. The inhibitory postsynaptic current was applied for 20 ms. The constant input into the neuron (ii) was chosen to obtain different baseline firing rates of approximately 10, 14 and 18 Hz.

The noise component (iii) was composed of two parts, a zero-mean, band-pass filtered (15 Hz to 35 Hz) and a zero-mean low-pass filtered (*<* 100 Hz) white noise [46, 47]. Thereby, the standard deviation of the noise input was scaled with respect to the mean drive such that the band-pass filtered component accounts for 80% of the total standard deviation [48]. To investigate the effect of noisy inputs in the investigated reflex scenario, we applied different noise levels by scaling the total standard deviation of the summed noise input to 0 %, 12.5 % and 25 % of the constant input current (ii). The input components (i) to (iii) were linearly summed and applied to the soma compartment, assuming they represent effective synaptic currents [28, 18]. To account for the delay caused by the nerve conduction velocity, a constant delay of 50 ms between stimulus application and membrane potential output was introduced.

### 4.5 Data Analysis

Motoneuron activity is described by means of discrete discharge times, i.e., a binary spike train. For the *in vivo* experiment, spike trains were reconstructed by the decomposition of the recorded HDsEMG [for details, see 7, 40, 41]. In the simulation, the spike trains were directly obtained from the trajectory of the soma membrane potential.

The firing behavior of motoneurons before and after stimulation was analyzed using the peristimulus frequencygram (PSF) method (Fig 2a). The PSF shows the instantaneous discharge rates of motoneurons relative to the stimulus time [20]. Significant perturbations in discharge rate compared to baseline activity after the stimulus are considered to be likely related to the stimulating event. With increasing delay between the stimulus and the event, the probability that changes in discharge rate may reflect other processes increases [49, 50, 51]. The characteristics of the PSF response are assumed to depend both on the sign as well as the magnitude of a (reflex-induced) postsynaptic potential [51]. Although other factors may play a role, PSF is recognized as a valid method to quantitatively estimate *in vivo* the strength of inhibitory or excitatory postsynaptic potentials in motoneurons [49, 50, 51].

The strength of reciprocal inhibition and the consecutive excitation was determined by computing the cumulative sum of the PSF (PSF-cusum, Fig 2b) [52]. The PSF-cusum allows the measurement of subtle but consistent changes in the instantaneous discharge frequency. It was obtained by cumulatively summing the difference between the average baseline frequency (*−*300 ms *≤* time *<* 0 ms) and the instantaneous frequency value of each discharge.

The largest absolute deflection of PSF-cusum from zero during the baseline was determined as the significance threshold (error box) for reflex responses (Fig 2b, dashed lines). Troughs and peaks exceeding the error box are considered to represent significant inhibition and excitation responses respectively. The response amplitude (inhibition or excitation) was defined as the difference between the onset and the plateau of the peak or trough in the PSF-cusum (Fig 2b, arrows). The onset and the knee point of the plateau were determined manually. The PSF-cusum was normalized by the number of delivered stimuli (i.e., trigger) to compare amplitudes across subjects and trials with different numbers of stimuli. Consequently, PSF-cusum is shown in units of Hz*/*trigger. It is worth noting that, in simulations where the membrane noise is not added, each deflection from zero is directly related to a change in the motoneuron’s activity. Recorded motor units were included in the analysis when at least 90 stimuli could be delivered. For simulated neurons always 200 stimuli were applied.

## Author Contributions

FN, LS, TK and UY conceptualized the study; LS developed the model and conducted the simulation and the data analysis. All authors contributed to the analysis and discussion of the results as well as to the writing of the manuscript.

## Acknowledgments

Laura Schmid was funded by Deutsche Forschungsgemeinschaft (DFG, German Research Foundation) under SPP 2311 (465243391). Thomas Klotz was supported by the ERC-AdG “qMOTION” (Grant agreement ID: 101055186). Francesco Negro was supported by the European Research Council Consolidator Grant INcEPTION (contract no. 101045605).

## Supplementary Figures

**Fig S 1.**
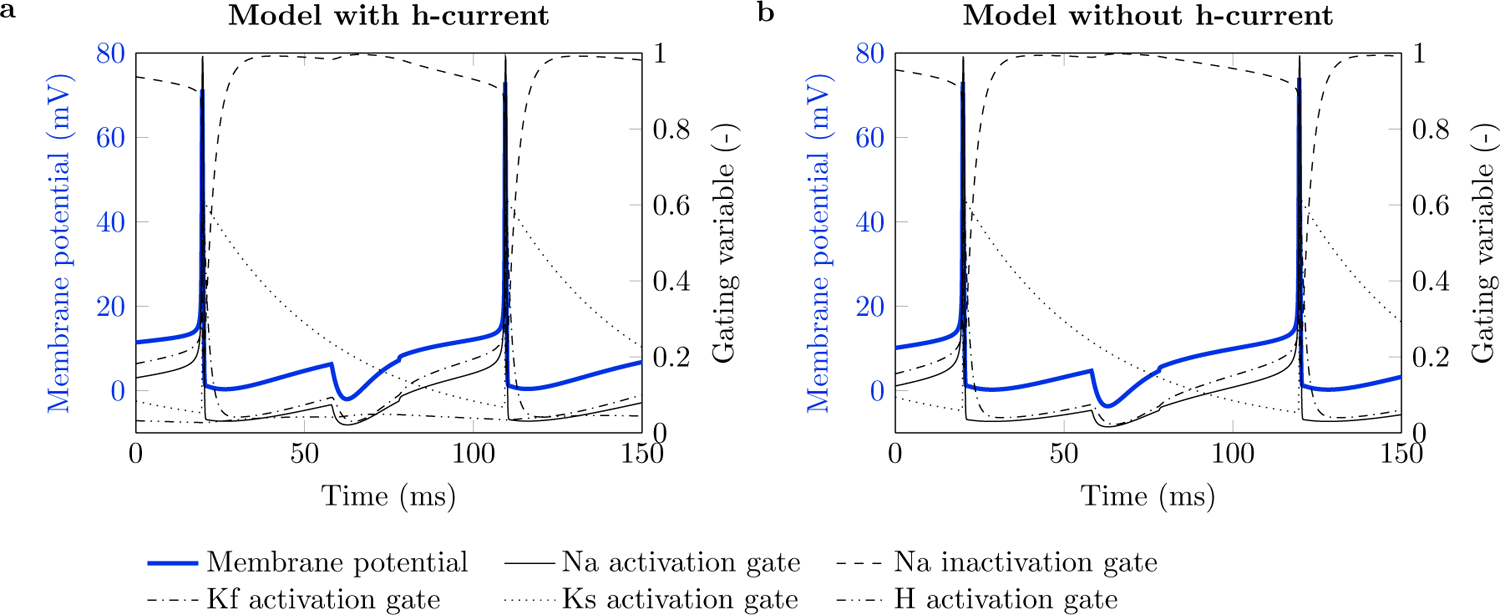
Motoneuron model gating variables. Motoneuron model gating variables for the model with (a) and without (b) h-current. The membrane potential trajectories (blue) correspond to the ones shown in blue in Fig 5. Gating variables (black) are defined as described in Negro and Farina [18] and Röhrle et al. [39].

**Fig S 2.**
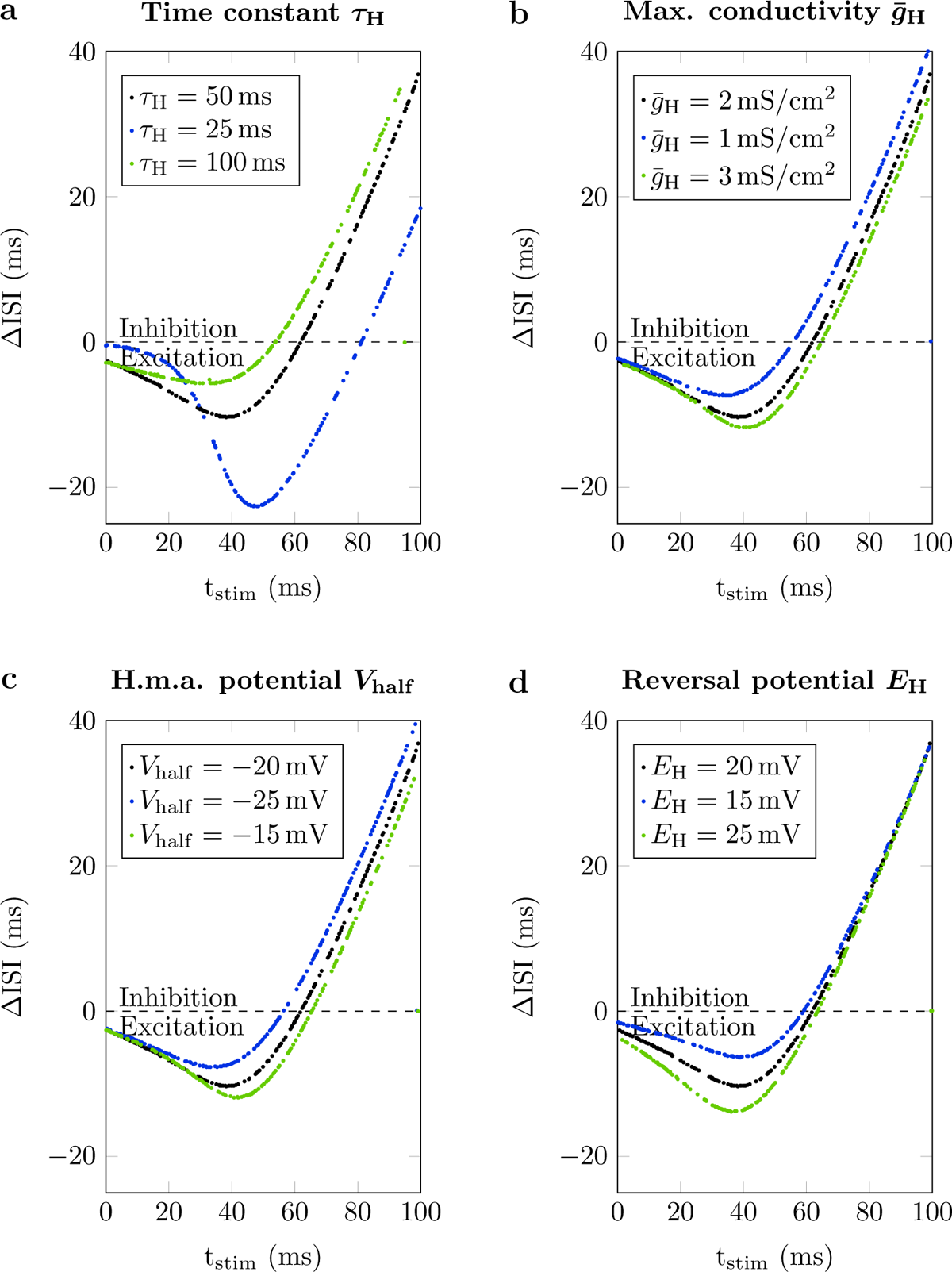
Stimulus time-dependent duration of interspike intervals for different parameterizations of the h-channel. Change of interspike interval duration (Δ ISI) over time of stimulus application with respect to the last discharge (t_stim_). Default parameters are shown in black. In the simulations all parameters were fixed except for time constant *τ*_H_ (a), maximum conductance of h-current *g*^-^_H_ (b), half maximum activation potential *V*_half_ (c) or reversal potential *E*_H_ (d). Baseline frequency 10 Hz, no noise, inhibitory stimulus amplitude *−*10 nA.

## Notes

### Competing Interest Statement

The authors have declared no competing interest.

